# Homotypic endoplasmic reticulum membrane tethering is critical for flavivirus replication

**DOI:** 10.1101/2025.11.26.690702

**Authors:** Jonathan Einterz Owen, Cheyanne Lee Bemis, Qingyi Wang, Ambarish C. Varadan, Jacob W. Vander Velden, Laura Andačić, Olus Uyar, Mansi Gupta, Christopher D. Scharer, Laurent Chatel-Chaix, Pietro Scaturro, Mehul S. Suthar, Christopher J. Neufeldt

## Abstract

Flaviviruses (genus *Orthoflavivirus*) are arthropod-borne viruses which cause approximately 400 million annual global infections in humans. Flavivirus infection requires cellular machinery to facilitate replication and spread. All known flaviviruses replicate in association with the host endoplasmic reticulum (ER), where genome replication is confined within virus-induced ER invaginations called viral replication organelles (vROs). Despite the central role of these structures during flavivirus infection, the mechanisms underlying vRO biogenesis remain undefined – particularly the membrane rearrangements required for their formation. In this work, we report a conserved role for a cellular ER remodeling protein, atlastin-2 (ATL2), in the organization of vROs within infected cells. Using confocal and electron microscopy, we show that ATL2 depletion leads to a reduction in vRO spatial distribution in flavivirus-infected cells. Changes in vRO distribution corresponded with a decrease in virus production and robust induction of innate immune responses. We also demonstrate that ATL2 accumulates in areas of vRO formation during flavivirus infection. Critically, mutational analysis showed that a tethering-competent but fusion-defective ATL2 mutant was sufficient to rescue DENV and ZIKV replication in ATL2-knockout cells. Finally, inhibition of ATL2 activity using synthetic peptides significantly reduced DENV replication in both immortalized and human primary cells, suggesting a possible avenue for targeting host ER functions to limit flavivirus replication. Taken together, these results show that membrane tethering plays a critical and conserved role in flavivirus infection, functioning to organize membranes for vRO biogenesis and limit cellular immune activation. Importantly, we provide evidence that ATL2-mediated membrane organization can be targeted to inhibit viral replication.

## Introduction

*Orthoflavivirus* (formerly *Flavivirus*) is a genus of arboviruses that contains numerous pathogens of significant consequence to human health, including West Nile virus (WNV), Zika virus (ZIKV), and dengue virus (DENV). These pathogens are responsible for an estimated 400 million human infections per year, with more than 80% of countries and half the world’s population at risk for infection^1^. The global prevalence of flaviviruses, combined with their high mutation rates, makes recurrent epidemics or novel emergence events likely^2,3^. This is exemplified by the recent emergence, or re-emergence, of flaviviruses such as WNV, yellow fever virus (YFV), Powassan virus (POWV), and ZIKV into new human populations, highlighting the potential for novel flaviviruses to become highly dangerous to human hosts^3–8^.

Given their relevance to human health and lack of commercially available antiviral therapies, there is a pressing demand for therapeutic interventions that treat both emerging and endemic flaviviruses^9^. Host-targeted antivirals, which disrupt conserved virus-host interactions, offer an attractive possibility for broad-spectrum drug development – particularly because the barriers to resistance are theoretically higher than those of direct-acting antivirals^10,11^. Therefore, identifying host pathways that are critical for a wide array of virus infections represents an important avenue for antiviral intervention.

Replication of the flavivirus positive-sense single-stranded RNA genome occurs in association with the endoplasmic reticulum (ER) of host cells. More specifically, flavivirus genome replication is presumed to occur inside spherical virus-induced alterations of the ER membrane called viral replication organelles (vROs)^12–14^. The formation of vROs is highly conserved during flavivirus infections, with remarkable similarity in membrane morphology observed across disparate flaviviruses^15^. A vRO consists of a membranous vesicle formed by invagination of the ER into its luminal space, with a diameter of 70-90 nm and a connection to the cytosol through a pore-like opening^13,16–19^. Compartmentalization of viral genome replication within vROs is thought to serve several purposes, including concentrating the necessary proteins and substrates for RNA replication, coordinating trafficking of newly synthesized viral RNA, and shielding viral replication intermediates from immune receptors^16,20–26^.

Morphological data and comparisons with other positive-sense single-stranded RNA viruses suggest that new copies of the viral genomic RNA exit the vRO through the pore-like opening and are then engaged in one of two distinct viral processes: translation by ribosomes to produce viral proteins and form new vROs, or packaging into assembling viral particles^27–32^. However, the molecular mechanisms of flavivirus vRO biogenesis and the regulation of viral RNA trafficking in infected cells remain unclear.

*De novo* vRO biogenesis and virion assembly typically occur on ER membranes proximal to sites of existing vROs^14,17,18,26^. The close connection between ER membranes during viral processes is thought to ensure efficient RNA movement while minimizing exposure of the viral genome to cytosolic RNA sensors and the RNA degradation machinery of the innate immune system^23,33,34^. However, the mechanisms that facilitated these membrane contacts and how this membrane organization specifically support infection remain undefined. In previous works, we and others implicated an ER-resident membrane remodeling protein called atlastin-2 (ATL2) in the replication of DENV and ZIKV^35,36^. ATL2 is a membrane fusion protein which uses GTP binding and hydrolysis to dimerize across opposing ER membranes, leading to membrane fusion and the formation of three-way junctions in the peripheral ER that help to maintain ER branching and homeostasis^37,38^. Using liposomes *in vitro*, ATL2 dimers have also been observed to tether juxtaposed membranes without fusing them, though the physiological role of tethering remains under investigation^39–42^.

In this study, we define a conserved role for ATL2 membrane remodeling during flavivirus infection. Our results indicate that ATL2 is critical for vRO distribution and organization during replication. We found that ATL2 accumulates at sites of vRO biogenesis, and that its membrane tethering activity was required to facilitate proper viral replication. We further linked the disruption of proper vRO formation in ATL2-depleted cells to increases in immune activation. Finally, we leveraged insights into ATL2 tethering function to design inhibitory peptides that impede ATL2 activity and limit virus infection. Our results establish ER-to-ER membrane tethering as a critical host process for the spatial organization of flavivirus vROs and demonstrate that this organization is important for viral evasion of cellular immunity. Additionally, our work provides a proof-of-concept for using mechanistic interrogations of host-virus interactions to identify potential therapeutic intervention targets.

## Methods

### Viruses and cell lines

Vero E6, HEK-293T, C6/36, and A549 cells were obtained from ATCC. Huh7 cells were obtained from the research group of Dr. Arash Grakoui (Emory University). Huh7-Lunet-T7 cells were obtained from the research group of Dr. Ralf Bartenschlager (University of Heidelberg). All cell lines were regularly tested for mycoplasma contamination; cell lines were authenticated by visual observations of cell morphology. All cells were cultured in Dulbecco’s modification of Eagle’s medium (DMEM, Corning) supplemented with 10% fetal bovine serum, 100 U/mL penicillin, 100 µg/mL streptomycin, and 1% non-essential amino acids (complete media). Stable A549 cell lines were cultured in complete media containing either 5 μg/mL blasticidin or 1 μg/mL puromycin. Huh7-Lunet-T7 cells were cultured in complete media containing 2 μg/mL zeocin.

Molecular clones of DENV genotype 2 (strain 16681), ZIKV (strain H/PF/2013), and the reporter *Renilla* luciferase (RLuc) DENV (strain 16681; DV-R2A) have been previously described and were a gracious gift of Ralf Bartenschlager^43–47^. Isolates of WHO reference strains for DENV serotypes DV1-WP75, DV3-H87, and DV4-H241, as well as CHIKV 181/25 and ONNV UgMP30, were generously provided by Dr. Matthew Collins (Emory University). For molecular clones, virus stocks were prepared as previously described by electroporation of Vero E6 cells with *in vitro* transcribed viral RNA; harvesting occurred 4-8 days after electroporation^45^. For amplification of isolates or stocks from molecular clones, VeroE6 or C6/36 cells were infected at an MOI of 0.01 and supernatants were harvested 2-8 days after infection, beginning when cells showed cytopathic effect.

Extracellular virus titers were determined by plaque-forming unit (PFU) assay in Vero E6 cells using a minimum essential medium (MEM, Gibco) overlay containing 1.0% carboxymethylcellulose (CMC). For DENV serotypes 1, 3, and 4, titers were determined using a focus-forming unit (FFU) assay; infected cells in MEM-1.0% CMC were fixed and stained for the viral NS3 protein and infection foci were counted using fluorescence microscopy^48^. Titers of the DV-R2A virus, which does not produce plaques on Vero E6 cells, were also calculated using a fluorescence-based approach. Briefly, cells were infected at 10-fold decreasing dilutions by DV-R2A or by a wildtype DENV with a known titer. The percentage of infected cells was determined by immunofluorescence staining for the viral NS3 protein (see below). A sigmoidal regression curve, using the percentage of infected cells caused by the wildtype virus as a standard, enabled quantitation of the R2A virus titer.

### ATL2 expression and depletion constructs

A list of all primers used for ATL2 cloning is provided in **Supp Table 1**. For all ATL2 expression constructs, the NEBuilder HiFi cloning system was used as per the manufacturer’s protocol (New England Biolabs). ATL2 mutants were amplified from full-length ATL2-containing plasmids in the pWPI backbone (a gift from Didier Trono, Addgene plasmid #12254); we previously introduced synonymous mutations in the ATL2 gene sequence to remove cryptic bacterial promoters and facilitate plasmid amplification in bacterial strains^35^. Sequences were transferred to the pWPI backbone using the NEBuilder HiFi DNA Assembly Master Mix (New England Biolabs) and subsequently transformed into competent DH5α *Escherichia coli* (Invitrogen).

Successful transformants were confirmed using Sanger sequencing (Azenta Life Sciences). Constructs for shRNA-mediated ATL2 depletion, as well as non-target (NT) shRNA control constructs, have been described previously^35^.

### Lentivirus production and transduction of cells

Delivery of expression plasmids for depletion and overexpression experiments was accomplished through transduction with lentiviruses. For production of lentiviral stocks, sub-confluent 293T cells were transfected with packaging plasmids pCMV-Gag-Pol (Addgene plasmid #22036) and pMD2-VSV-G (Addgene plasmid #12259), kind gifts from Didier Trono, as well as pWPI insert plasmids. Two days post-transfection, lentivirus-containing media was collected and filtered through 0.45 µm PVDF syringe filters (MilliporeSigma). Lentiviruses were titrated by SYBR Green I-based real-time PCR-enhanced reverse transcriptase (SG-PERT) assay, using the iTaq Universal SYBR Green Supermix (Bio-Rad Laboratories)^49,50^. Lentivirus stocks were compared to a standard curve derived from a sample with a known RNA concentration in order to determine the titer for each stock.

Transductions were performed using an MOI of 5 in the presence of 4 µg/mL polybrene, with the following exceptions. All experiments utilizing glass coverslips for immunofluorescence (see below) did not contain polybrene due to an unknown, previously unobserved, and inexplicable adverse reaction between polybrene-containing media and our glass coverslips, which resulted in cell death.

Transductions of Huh7 or Huh7-Lunet cells were performed at an MOI of 10 owing to reduced transduction efficiencies in these cells. Stable cell lines were produced by transduction of target cells with lentivirus at an MOI of 1, followed by selection with media containing either 5 μg/mL blasticidin or 1 μg/mL puromycin, depending upon the resistance marker of the lentiviral vector. Viability of transduced cells was evaluated using the CellTiter-Blue Cell Viability Assay (Promega) by measuring fluorescence (excitation 560 nm, emission 590 nm) with a BioTek Synergy H1 multimode microplate reader (Agilent Technologies) according to the manufacturer’s instructions.

### Reverse transcription - quantitative PCR (RT-qPCR)

Cells were washed once with PBS, followed by lysis with RNA Lysis Buffer (Zymo). Total cellular RNA was isolated using a Quick-RNA Miniprep RNA extraction kit (Zymo) according to the manufacturer’s protocol. Synthesis of cDNA was accomplished using a high-capacity cDNA reverse transcription kit (Applied Biosystems) according to the manufacturer’s specification. cDNA was used for qPCR analysis after being diluted 1:10, using specific primers and the iTaq Universal SYBR Green Supermix (Bio-Rad).

Primers for qPCR were designed using Primer3 software; primer sequences are supplied in **Supp Table 2**. To obtain the relative abundance of specific RNAs from each sample, cycle threshold (Ct) values for cDNAs of interest were normalized to those of hypoxanthine phosphoribosyltransferase 1 (HPRT). All qPCR runs were performed on a C1000 Dx Thermal Cycler with a CFX96 Optical Reaction Module (Bio-Rad).

### Western blotting

Cells were lysed using 2X Laemmli buffer, followed by sonication and denaturation at 95 °C for 5 minutes. ∼10 µg of total protein from each sample was resolved with a 10% SDS-PAGE gel and transferred to a nitrocellulose membrane. Membranes were blocked with PBS-T (PBS [pH 7.4] with 0.1% Tween-20) containing 5% skim milk for 1 h at room temperature. Primary antibodies were incubated in PBS-T containing 2% skim milk overnight at 4°C. Membranes were washed with PBS-T and incubated for 1 h with the appropriate HRP-conjugated secondary antibodies. For a list of primary and secondary antibodies used, see **Supp Table 3**. Following secondary incubation, membranes were again washed with PBS-T and Clarity Western ECL Substrate (Bio-Rad) was applied. Membranes were imaged using the ChemiDoc MP Imaging System (Bio-Rad). Images were cropped and analyzed using Fiji software^51^.

### Immunofluorescence (IF)

A549 cells grown on 12 mm glass cover slips (Azer Scientific) or in µCLEAR black-walled 96-well plates (Greiner BioOne) were washed 1x with PBS, fixed with 4% paraformaldehyde at room temperature for 20 min, washed 3x with PBS, and permeabilized in 0.2% Triton X-100 for 2 min at room temperature. Following another wash step (3x with PBS), samples were blocked in 2.5% bovine serum albumin (BSA) or 2.5% skim milk in PBS-0.01% Tween-20 for 1 h at room temperature, then incubated with the indicated primary antibodies (diluted in blocking buffer) at 4°C overnight (for a list of primary and secondary antibodies used, see **Supp Table 3**). Samples were then washed 6 times with PBS-0.01% Tween-20 (3x10 sec, 3x10 min) and incubated with secondary antibodies in PBST for 45 min at room temperature. Following another 6 washes (3x10 sec, 3x10 min) coverslips were mounted onto microscope slides using DAPI Fluoromount-G mounting media (SouthernBiotech). For 96-well plates, DAPI was added at 2.86 µM for 20 min prior to secondary washes.

Epifluorescence images were obtained using an Axio Observer Z1 microscope with an Axiocam 506 monochromatic camera (Zeiss) or a BioTek Lionheart FX automated widefield microscope (Agilent). Confocal images were obtained with a Leica Stellaris 8 Inverted Confocal Microscope (Leica Microsystems) with a white light laser and both HyD-S and Power HyD-X detectors. Image analysis to determine percent infection, fluorescent signal intensity, and fluorescent signal distribution was performed using CellProfiler^52^. Pearson’s correlation coefficients were calculated using the Coloc2 plugin for Fiji. Adobe Photoshop and Illustrator software packages were used to assemble images into figures.

### Proximity ligation assays (PLA)

Coverslips were washed twice with PBS, fixed with 4% paraformaldehyde for 20 minutes at room temperature, and permeabilized with 0.2% Triton X-100 for 20 minutes. PLA was performed using the Duolink PLA Kit (Millipore-Sigma) according to the manufacturer’s instructions and as previously described^48,53^. Briefly, cells were blocked in the kit buffer for 1 hour at 37°C, then incubated with anti-HA and anti-NS5 primary antibodies for 2 hours at room temperature. For a list of antibodies used, see **Supp Table 3**. After washing, coverslips were incubated with PLUS and MINUS PLA probes for 1 hour at 37°C, followed by ligation (30 minutes) and amplification (100 minutes) at 37°C. Cells were stained with DAPI (Life Technologies), washed, and mounted using Fluoromount-G (Southern Biotechnology Associates). Images were acquired using an LSM780 confocal microscope (Zeiss) at the INRS Confocal Microscopy Core Facility.

PLA dot quantification was performed with Fiji software. Signals >0.02 μm² with circularity between 0.02 and 1.00 were counted per cell. Identical acquisition and analysis settings were applied across all experiments.

### Electron microscopy (EM)

Cells grown in 24-well plates were washed with pre-warmed PBS, then fixed for 30 min by incubation with pre-warmed 1.25% glutaraldehyde in 0.2 M HEPES-buffered H_2_O. After 3 washes with 0.2 M HEPES-buffered H_2_O, cells were incubated with 2% osmium tetroxide/0.2 M HEPES-buffered PBS for 40 min on ice. Cells were washed with water 3 times and treated with 0.5% uranyl acetate for 30 min. After a 30-min wash with water, cells were progressively dehydrated with increasing concentrations of ethanol (40% to 100%), followed by a wash with propylene oxide and the addition of Epon/araldite resin (Araldite 502/Embed 812 kit; Electron Microscopy Sciences), which was then polymerized at 60°C for 72 h. Embedded cells were sectioned into 70-nm-thick slices by using an Ultracut UCT microtome (Leica) and a diamond knife (Diatome). After counterstaining with 3% uranyl acetate in 70% methanol for 5 min and 2% lead citrate in water for 2 min, cells were imaged with a JEOL JEM-1400 120kV LaB6 transmission electron microscope (TEM) with a Gatan US1000 CCD camera (Jeol Ltd., Tokyo, Japan). Whole-cell overview images at 4000X or 8000X magnification were obtained using SerialEM^54^. Quantification of vRO number and location was performed manually in Fiji. Distribution of vROs was quantified using a custom statistical pipeline developed using R and ChatGPT (OpenAI, 2025)^55–59^. For statistical analysis of vRO distribution, only those cells with greater than 20 vROs were used. For tomography images, 200-nm-thick sections were prepared, and 10 nm-diameter protein A-gold was added to both sides of the grid. Grids were placed in a high-tilt holder, and digital images were recoded as single-axis tilt series over a −60° to +60° tilt range (increment 1°).

Tomograms were reconstructed using the IMOD software package (https://bio3d.colorado.edu/imod) and segmented manually using Dragonfly 3D World software (Comet Technologies Canada)^60^.

### Correlative light and electron microscopy with immunofluorescence (immuno-CLEM)

Immuno-CLEM was performed as previously described^61,62^. Briefly, A549 cells expressing HA-tagged ATL2 were seeded into glass-bottom cell culture dishes containing etched gridded coverslips (MatTek Life Sciences) and infected with DENV or ZIKV (MOI = 5). After 48 h, cells were fixed in 4% PFA, 0.05% glutaraldehyde, and 0.2 M HEPES in H_2_O for 10 min at room temperature. Cells were washed 2 times with 4% PFA and 0.2 M HEPES in H_2_O, then fixed in this solution for an additional 30 minutes. After 3 five-minute washes with PBS, cells were incubated with blocking solution (50 mM NH_4_Cl, 0.1% saponin, 1% BSA in PBS) for 30 min. Cells were then incubated with primary antibodies diluted in blocking solution overnight at 4°C (for list of antibodies see **Supp Table 3**). The next day, following 6 two-minutes washes with PBS, secondary antibody diluted in blocking solution was applied for 1 hr at room temperature. DAPI was then added at 2.86 µM for 15 minutes. Following 6 more two-minute washes with PBS, cells were imaged using a Leica Stellaris 8 Inverted Confocal Microscope (Leica Microsystems). Z-stacks were obtained and the position of cells of interest on the etched grids was recorded using transmitted light (brightfield). Cells were then fixed and processed for EM as described above. Areas of interest for EM imaging were identified using the MatTek etched grid, visible in the Epon. EM samples were visualized with a JEOL JEM-1400 TEM. The shape of each cell’s nucleus (as indicated by DAPI signal area) was used to correlate and overlay EM images with the appropriate IF Z-slice; alignment was performed using ImageJ and Adobe Photoshop software packages.

### Plasmid-induced replication organelle (pIRO) polyprotein expression system

Huh7-Lunet-T7 cells, which stably express a cytosolic T7 RNA polymerase, were transduced with lentiviruses (MOI 5) containing either NT shRNA or ATL2 shRNA sequences and seeded into 6-well plates. After 2 days, transduced cells were seeded into 24-well plates at 30,000 cells/well. 24 hr later, cells were transfected with either the DENV or ZIKV pIRO construct described previously, using TransIT transfection reagent (Mirus Bio)^63,64^. 18 hr after transfection for DENV pIRO, or 16 hr after transfection for ZIKV pIRO, cells were processed for EM, IF or Western blot as described above. IF was used to evaluate transfection efficiency, as determined by the percentage of cells staining positive for the viral NS3 protein; Western blots were used to verify protein expression level. EM quantification of vROs was performed by systematically surveying cells and evaluating for the presence of vROs, followed by imaging at 8000X magnification. Only cells with >4 vROs were considered positive. For each condition, >50 cells were surveyed over 3 biological replicas. All observed vROs were imaged, and vRO diameters were determined using Fiji by measuring the distance across two axes and averaging.

### Bulk RNA sequencing (RNA-seq) analysis

A549 cells were transduced with constructs expressing ATL2 shRNA or NT shRNA for 4 days. Four biological replicates were used for each sample. Cells were lysed with RLT buffer (Qiagen) and RNA-seq libraries were prepared at the DLS HudsonAlpha using the NEB Ultra II RNA Library Preparation kit. Libraries were sequenced on a Novaseq X-plus PE100 run. Raw fastq reads were mapped to the hg38 human genome using STAR (v2.7.6a) with the Gencode V27 reference transcriptome^65,66^. Duplicate reads were removed from downstream analysis using the PICARD markduplicates (v2.23.8) function (http://broadinstitute.github.io/picard/).

Reads mapping to exons for all unique ENTREZ genes were compiled and normalized using GenomicRanges (v1.38.0) and all genes expressed at 3 or more reads per million in all samples of any one biological group were considered expressed^67^. Differential expression analysis was performed with DESeq2 (v1.42.1) and genes with an FDR < 0.05 and absolute log2 fold-change ≥1 were considered significant^68^.

### Proteomic analysis

A549 cells expressing lentivirus-delivered ATL2-targeted or NT shRNA for 4 days were scraped and pelleted at 100xg for 5 min, followed by freezing at -80°C. Thawed cell pellets were lysed in 250 µL of lysis buffer (10 mM Tris pH 7.0, 150 mM NaCl, 4% sodium dodecyl sulphate [SDS]) containing protease and phosphatase inhibitors (cOmplete and PhosStop, Roche), then sonicated. Clarified protein lysates were precipitated with acetone twice, and 50 µg of normalized protein mixtures was resuspended and denatured in 40 µL U/T buffer (6 M urea/2 M thiourea in 10 mM HEPES, pH 8.0). Proteins were reduced and alkylated in 10 mM DTT and 55 mM iodoacetamide, followed by digestion using 1 µg of LysC (FUJIFILM Wako Chemicals) and trypsin (Promega) in ABC buffer (50 mM NH_4_HCO_3_ in water, pH 8.0) overnight at 25°C, 800 rpm. After digestion, peptides were purified on stage tips with 3 layers of C18 Empore filter discs (3M) as previously described^69^.

Samples were analyzed on a Vanquish Neo LC system (Thermo Scientific) coupled to an Orbitrap Exploris 480 mass spectrometer (Thermo Scientific) equipped with a Nanospray Flex source (Thermo Scientific). Peptides were injected into an Acclaim PepMap 100 trap column (2 cm x 75 μm, 3 μm C18; Thermo Fisher Scientific) and next separated on a 60 cm x 75 μm column (1.7 µm C18 beads UHPLC column; Aurora Ultimate) with a packed emitter tip (Ion Opticks), at a constant flow rate of 300nl x min-1 over 120 min linear gradient. The column temperature was maintained at 45 °C using an integrated column oven (Sonation GmbH). The column was equilibrated using 3 column volumes before loading samples in 96% buffer A (99.9% Milli-Q water, 0.1% formic acid (FA)) / 4% buffer B [99.9% ACN, 0.1% FA]). Samples were separated using a linear gradient from 10 to 31% buffer B over 62 min before ramping up to 43% (33 min), 100% (1 min) and sustained for 10 min, followed by reduction to 4% buffer B (8 min). The Orbitrap Exploris 480 was operated in positive ion mode, with a positive ion voltage of 2000 V in data-dependent acquisition mode (DDA) using the Thermo Xcalibur software (v. 4.5.474.0). DDA analysis was performed with a cycle time of 1.5 sec.

Survey scans were acquired at 120,000 resolution, with a full scan range of 350- 1400 m/z, an automatic gain control (AGC) target of 300% and a maximum ion injection time of 25 ms, intensity threshold of 5 x 103, 2–6 charge state, dynamic exclusion of 90 sec, mass tolerance of 10 ppm. The selected precursor ions were isolated in a window of 1.6 m/z, fragmented by a higher-energy collisional dissociation (HCD) of 30.

Fragment scans were performed at 15,000X resolution with an Xcalibur-automated maximum injection time and standard AGC target.

### Raw mass spectrometry data processing and analysis

Raw MS data were processed with the MaxQuant software (v. 2.2.0.0) using the built-in label-free quantitation algorithm and Andromeda search engine^70^. The search was performed against the *Homo sapiens* proteome (UniprotKB release UP000005640; Taxon ID 9606) containing forward and reverse sequences. Additionally, the intensity-based absolute quantification (iBAQ) algorithm and “match between runs” option were used. In MaxQuant, carbamidomethylation was set as fixed and methionine oxidation and N-acetylation as variable modifications. Initial search peptide tolerance was set at 20 p.p.m. and the main search was set at 4.5 p.p.m. Experiment type was set as data-dependent acquisition with no modification to the default settings. Search results were filtered with a false discovery rate of 0.01 for peptide and protein identification. The Perseus software (v. 1.6.15.0) was used to further process the affinity-purification datasets. Protein tables were filtered to eliminate identifications from the reverse database and common contaminants. In the subsequent MS data analysis, only proteins identified on the basis of at least one peptide and a minimum of three quantitation events in at least one experimental group were considered. The MaxLFQ protein intensity values were median-normalized and log2-transformed, and missing values were filled by imputation with random numbers drawn from a normal distribution calculated for each sample^71^. Knockdown-specific proteome changes were determined by two-sided Student’s Test (S0=1) with permutation-based false discovery rate statistics (250 permutations, FDR threshold 0.05) or using a log2 (fold change) ≥ 2 cutoff. Results were plotted as scatter plots using Perseus^71^.

### *Renilla* luciferase assay

The expression of *Renilla* luciferase (RLuc) in cells infected with the DV-R2A reporter virus was used as a surrogate measurement of intracellular viral replication, as described previously^45,72^. Briefly, cells plated in white-walled µCLEAR 96-well plates (Greiner BioOne) and infected with DV-R2A reporter virus for 48 hr were lysed in 35 µL of RLuc lysis buffer (0.1% Triton X-100, 25 mM glycine-glycine [pH 7.8], 15 mM MgSO_4_, 4 mM EGTA, 10% glycerol, 1 mM DTT) and freeze-thawed at -80°C. RLuc activity was measured by adding 100 μl of Rluc assay buffer (15 mM potassium phosphate [pH 7.8], 25 mM glycine-glycine [pH 7.8], 15 mM MgSO_4_, 4 mM EGTA, 1.43 μM coelenterazine) to 15 μl of cell lysate and measuring luminescence using a BioTek Synergy H1 multimode microplate reader.

### Generation of ATL2-KO cells and ATL2-KO rescue experiments

The A549 ATL2 knockout (ATL2-KO) cells used in this work were briefly described previously and were created in the lab of Ralf Bartenschlager^35^. To ensure complete knockout of ATL2 while avoiding clonal effects, we created single cell clones by plating ATL2-KO cells at a limiting dilution to allow for the growth of single-cell colonies. Individual colonies were tested for ATL2 KO by Western blot (**Supp Fig 5A**), and 5 colonies which displayed complete ATL2 KO across 13 passages were pooled to create a new population of ATL2-KO pooled cells. In parallel, control cells with a nontarget guide sequence were produced using the same method; unaffected expression of ATL2 in control single-cell colonies was also confirmed by Western blot (**Supp Fig 5A**) prior to pooling 5 colonies, creating control pooled cells.

For ATL2-KO rescue experiments with DV-R2A, ATL2-KO pooled cells and control pooled cells were plated at 8,500 cells per well into 96-well plates and transduced with the specified lentiviruses at an MOI of 5. 24 hr after transduction, cells were infected with DV-R2A at an MOI of 1 for 48 h, with media replacement occurring at 24 hr post-infection. Viral replication was determined by RLuc assay (see above).

Individual experiments contained quadruplicate replicates, and results report all replicates from two independent experiments. For rescue experiments with WT ZIKV, cells were seeded at 40,000 cells per well into 24-well plates and transduced with specified lentiviruses at an MOI of 5. 24 hr after transduction, cells were infected with ZIKV at an MOI of 1 for 48 h, with media replacement occurring at 24 hr post-infection. Supernatants were collected, then cells were washed once with PBS and lysed with RNA lysis buffer. Viral replication was determined by qPCR and plaque assay (see above). Individual experiments contained two replicates, and results report all replicates from two independent experiments.

### Peptide treatment of infection

A549 cells were seeded at 7,000 cells per well into 96-well plates. The next day, ATL2 and control peptides (Peptide 2.0 Inc.) and DENV (MOI 1) were added to cells simultaneously. For a list of peptide sequences, see **Supp Table 4**. Peptides were applied at 200, 100, 50, or 25 µM concentrations. Media was changed after 1 h, with fresh peptide added (at the same concentrations). Cells were incubated for 48 h, at which point supernatants were harvested for plaque assay and cells were fixed for an IF-based assay to quantify infection levels (as described above). Results in all cases represent a minimum of two replicates from three independent experiments. Viability of peptide-treated cells was evaluated using the CellTiter-Blue Cell Viability Assay (Promega) according to the manufacturer’s instructions.

For peptide experiments in primary human monocyte-derived dendritic cells, MoDCs were isolated as previously described^73–75^. Briefly, whole blood was drawn from three anonymized donors and deposited into Vacutainer CPT tubes (BD Biosciences). Monocyte separation proceeded according to the manufacturer’s instructions. CD14+ cells were isolated using the MojoSort Human CD14 Selection Kit (BioLegend) as per the manufacturer’s instructions. These cells were resuspended in RPMI 1640 (Corning) supplemented with 10% FBS, 2 mM L-glutamine, 1 mM HEPES, 1 mM sodium pyruvate, 1% non-essential amino acids, 100 U/mL penicillin, and 100 µg/mL streptomycin (complete RPMI). 100 ng/mL of the cytokines granulocyte-macrophage colony-stimulating factor (GM-CSF) and interleukin-4 (IL-4) were also added to the media, and cells were plated into 6-well plates. One day after plating, cells were fed with fresh complete RPMI and cytokines. Five days after plating, non-adherent cells (differentiated moDCs) were collected and resuspended in complete RPMI, then plated at 100,000 cells/well into V-bottom 96-well plates. Cells were treated with the indicated peptide and infected with DENV or ZIKV (MOI 2), as above. Supernatants were harvested from each well at 24 and 48 hr post-infection. At 48 hr post-infection, cells were lysed for total RNA isolation. Viral titers were determined by plaque assay and viral RNA levels by RT-qPCR using the methods described above. Results represent duplicate samples from three different donors. Viability of treated cells was evaluated using the CellTiter-Blue Cell Viability Assay.

## Data availability

The mass spectrometry-based proteomics data have been deposited at the ProteomeXchange Consortium (http://proteomecentral.proteomexchange.org) via the PRIDE partner repository with the following dataset identifier: PXD068694. {Reviewer’s login details: Project accession: PXD068694, Token: uHlf7qX8hJVG.}. The bulk RNA sequencing data have been deposited at the GEO repository (accession: GSE309949).

## Ethics statement

Human peripheral blood mononuclear cells (PMBCs) were obtained from healthy donors in accordance with the Emory University Institutional Review Board according to IRB protocol IRB00045821.

## Results

### ATL2 depletion alters the intracellular distribution of viral replication sites for diverse flaviviruses

We first aimed to determine if the organization of membrane replication sites by the host GTPase ATL2 was conserved among flaviviruses and for other positive-strand RNA viruses^35^. The reported cellular function of ATL2 as an ER-shaping protein led us to hypothesize that ATL2 depletion would broadly negatively impact flavivirus infections while minimally impacting alphaviruses, which primarily use the plasma membrane or endosomal/lysosomal membranes for replication^76,77^. To test this, ATL2 was depleted in human A549 cells using lentivirus-transduced ATL2-targeted short hairpin RNA (shRNA), and cells were infected with representative flaviviruses or alphaviruses. Cells were monitored for cell viability, intracellular genome replication, and viral titers. Results from qPCR and plaque assays showed a significant decrease in viral titers and intracellular viral genome levels for all flaviviruses tested – including all four serotypes of DENV as well as ZIKV, WNV, and POWV – in ATL2 knockdown (ATL2 KD) cells, as compared to cells transduced with a nontarget (NT) shRNA construct (**Fig 1A-B and Supp Fig 1A-B**). Conversely, we observed that ATL2 KD did not significantly reduce genome replication in cells infected with the alphaviruses O’nyong’nyong virus (ONNV) and chikungunya virus (CHIKV). Importantly, cell viability analysis showed that depletion of ATL2 does not have a detrimental effect on cell growth that would generally limit virus replication and assembly (**Supp Fig 1C**). Together, these results indicate that the requirement for ATL2 is conserved for flaviviruses that replicate on ER membranes, and that ATL2 does not have a significant role in alphavirus replication, which occurs primarily at the plasma membrane.

**Fig 1.**
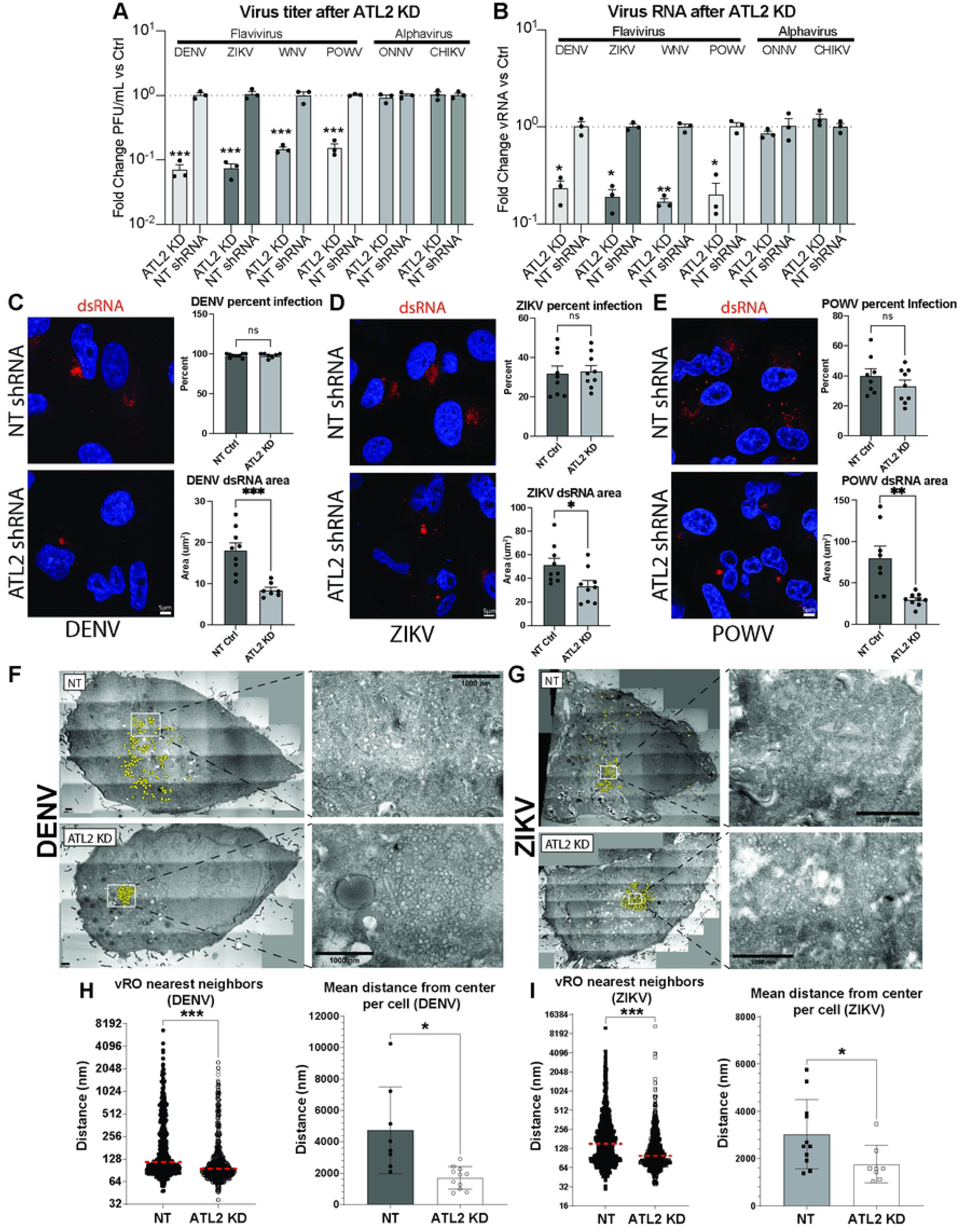
Impact of ATL2 depletion on flavivirus replication. A549 cells were transduced with the indicated shRNA constructs for 48 hr followed by infection with the indicated virus. **A-B**, Transduced cells were infected with DENV2 (strain 16681), ZIKV (strain H/PF/2013), POWV (strain Spooner), ONNV (strain UgMP30) or CHIKV (strain 181/25) for 48 hr followed by evaluating virus titer (**A**) and intracellular viral RNA (**B**). **A,** Graphs showing the mean fold change in PFU/ml relative to the non-targeting (NT) shRNA cells. Error bars show standard error of the mean (SEM); *n* = 3 biological replicates. **B**, Graphs showing the mean fold change in viral RNA levels relative to the non-targeting shRNA cells. Error bars show SEM; *n* = 3 biological replicates. For **A-B**, statistical significance was determined using one-way ANOVA with Dunnett’s multiple comparison analysis. *P < 0.05, **P < 0.01, ***P < 0.001. **C-E**, Cells were fixed with PFA and viewed by immunofluorescence microscopy using antibodies of given specificities. Images are representative of 3 independent experiments. Graphs show the average percent infection (top graph) and the average area per cell occupied by dsRNA signal (bottom graph). Scale bars, 5 μm. **F-I**, NT control and ATL2 shRNA transduced cells were infected with DENV for 48 hr (panels **F, H**) or ZIKV for 24 hr (panels **G, I**) for 24 hr, and viewed by TEM. **F-G,** Representative TEM images of DENV- or ZIKV-infected NT or ATL2 KD cells. vROs (yellow dots) were identified manually. Insets display magnified views of the boxed area. **H-I**, Centrographic statistics were used to quantify vRO distributions in NT or ATL2KD cells infected with either DENV or ZIKV. Left graph in each panel shows the distance between all vROs and their nearest neighbor (red line indicates median value). Right graph in each panel shows the mean distance from center of the vRO distribution for each cell. Statistical significance was determined using a Mann-Whitney test for nearest neighbor data and Welch’s t-test for mean distance from center data. N≥250 vROs per condition. For all graphs, *P < 0.05, ***P < 0.001.

We next evaluated the effect of ATL2 depletion on the localization of flavivirus replication sites, using viral double-stranded RNA (dsRNA, a viral replication intermediate) as a marker for viral genome replication. ATL2-depleted cells infected with either DENV, ZIKV, or POWV were stained for dsRNA and imaged using confocal microscopy. In flavivirus-infected ATL2 KD cells, we observed that dsRNA was primarily localized to a single punctum in the perinuclear region, exhibiting a reduced total area per cell compared to cells expressing non-target (NT) shRNA (**Fig 1C-E**). These results show that ATL2 depletion affects the spatial organization of replication sites across three diverse flaviviruses, suggesting that the mechanism of ATL2 function during flavivirus infection is conserved.

Since dsRNA forms as a replication intermediate inside vROs, the change in distribution of dsRNA indicates that vRO organization is likely altered in ATL2-depleted cells^78^. To determine the impact of ATL2 KD on vRO organization, we compared vRO distributions in ATL2 KD and control cells using transmission electron microscopy (TEM) (**Fig 1F-I**). Tiled images of NT or ATL2 KD cells were used to manually identify vRO locations (**Fig 1F-G**). Centrographic statistics were used to describe vRO distributions in each cell. For both DENV and ZIKV, ATL2 KD significantly decreased both the distance between a given vRO and its nearest neighbor, as well as the mean distance from center for a given cell’s vRO distribution, as compared to NT cells (**Fig 1H-I**). Decreases were also noted in the average nearest neighbor distance per cell (i.e. the average distance between a vRO and its closest neighbor, within each cell) and standard distance per cell (i.e. the square root of the sum of the squared distances from each vRO to the mean center of the vRO distribution for each cell) (**Supp Fig 1D-I**).

Taken together, these results demonstrate that ATL2 depletion alters the spatial organization of vROs for multiple flavivirus infections, causing vROs to form closer together and concentrate around the center of their distribution.

### ATL2 depletion disrupts vRO formation without disrupting protein production

To further evaluate the role of ATL2 in vRO biogenesis, we next measured the size of vROs from ATL2-depleted cells. We previously reported that ATL2 depletion altered the diameter of DENV vROs. However, these data were obtained from EM thin sections which cannot accurately depict three-dimensional (3D) morphology. To determine if ATL2 KD resulted in an overall decrease in the 3D volume of vROs, we used electron tomography on DENV-infected NT or ATL2 KD cells (**Fig 2A-C**).

**Fig 2.**
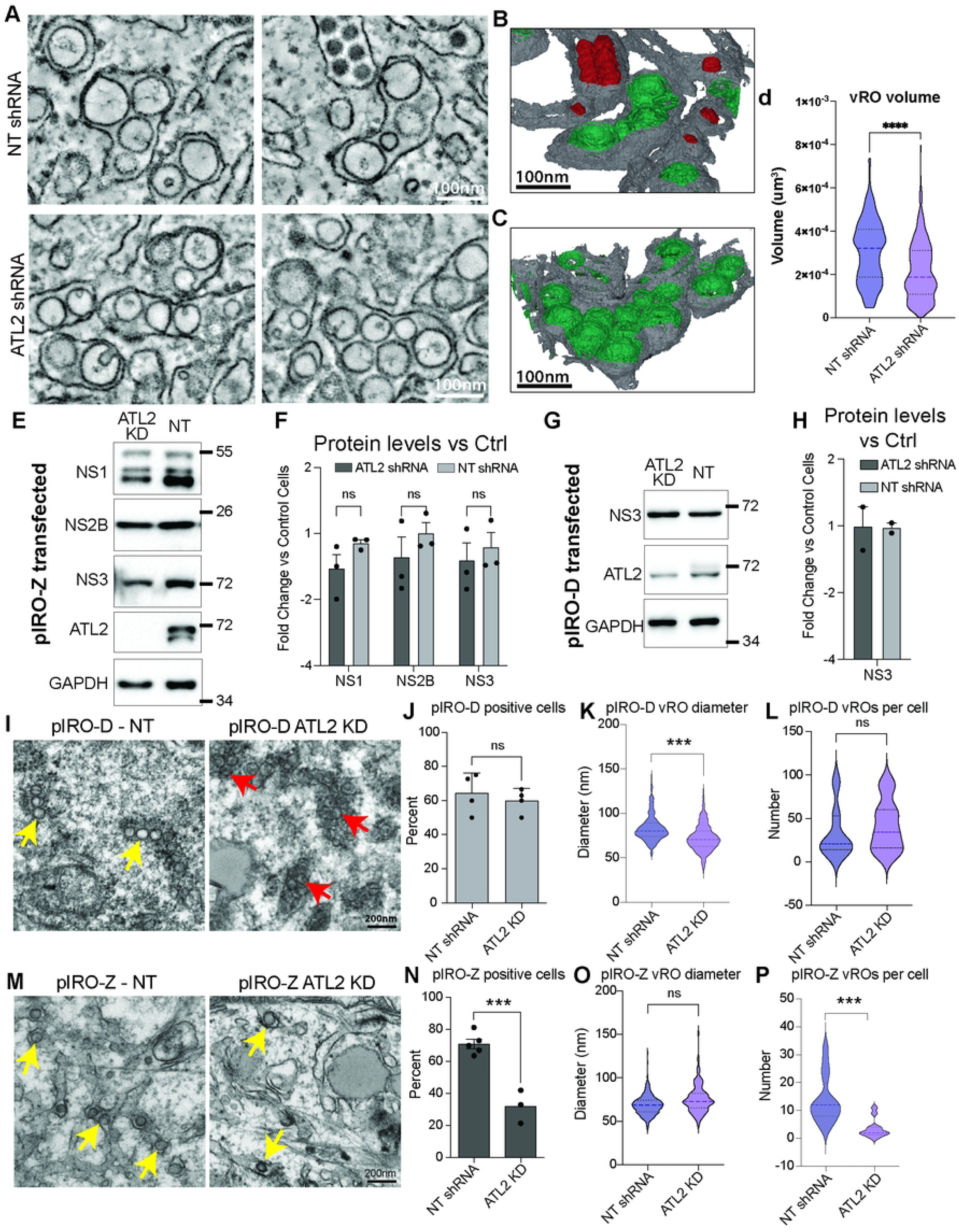
ATL2 depletion does not impact cellular or viral translation. **a-d**, Cells were transduced with constructs encoding for shRNA directed against ATL2, or a NT shRNA. 48 hr after transduction, cells were infected with DENV for 48 hr and analyzed by electron tomography. **A,** Representative single slices of reconstructed tomograms. Two images are shown for NT shRNA and ATL2 shRNA. Scale bars, 100nm. **B-C,** Reconstruction and segmentation of tomograms for area shown in **A**. **D,** Graph shows the average total volume of vROs from reconstructed tomograms. N≥250 vROs per condition. For all graphs, *P < 0.05, ***P < 0.001. **E-P,** Lunet-T7 cells were transduced with ATL2 or NT shRNA for 48 hr, followed by transfection with either pIRO-Z (**E-F** and **M-P**) or pIRO-D (**G-H** and **I-L**) constructs. **E-H**, Cell were lysed and viral protein levels evaluated using Western blotting with the indicated antibodies. **F, H,** Graphs show the average fold change in protein levels between ATL2-KD and NT control cells. Statistical significance was determined using one-way ANOVA with Dunnett’s multiple comparison analysis. *n* = 3 biological replicates for (**F**) and 2 biological replicates for (**H**). **I, M,** Representative EM images for pIRO-Z- and pIRO-D-expressing cells. **J-L** and **M-P**, Graphs show the average percentage of cells that contain vROs (**J, N**), the average vRO diameter (**K, O**) and the average number of vROs per positive cell (**L, P**). *n >* 40 cells for >3 biological replicates, error bars represent SEM. For all graphs ***P < 0.001, as determined by two-tailed T-test.

Evaluation of 3D-reconstructed vROs from NT cells showed an average volume of 3.1x10^-4^ μm^3^, consistent with previous reports for DENV infection (**Fig 2D**)^17^. In ATL2 KD cells, vROs had a significantly lower total volume of 2.16x10^-4^ μm^3^, indicating that ATL2 may play a direct role in vRO formation.

The formation of flavivirus vROs requires the accumulation of several viral non-structural proteins as well as specific RNA structures in the viral genome^64^. Therefore, it is possible that the observed changes in vRO organization following ATL2 depletion could result from decreases in viral protein translation or polyprotein processing. To determine if ATL2 depletion leads to global changes in cellular translation, we first compared total mRNA and protein levels between uninfected NT and ATL2 KD cells by integrating bulk RNA-seq and quantitative proteomics (**Supp Fig 2A-E**). Comparisons of differentially regulated transcripts and proteins identified a group of 103 host targets consistently upregulated at both the mRNA transcript and protein level, as well as 115 host targets consistently downregulated at both the transcriptional and translational levels, in ATL2 KD cells (**Supp Fig 2A**). Among the genes significantly upregulated following ATL2 depletion, we noted a slight functional enrichment for factors involved in inflammatory responses and TNF signaling (**Supp Fig 2A and 2D**). To specifically investigate possible selective effects of ATL2 on cellular translation, we additionally analyzed host proteins displaying altered abundance in the absence of significant changes at the mRNA level. This approach revealed only a small number of up- or down-regulated proteins in ATL2 KD cells when compared to NT controls, suggesting a lack of translation shut-off in ATL-2 KD cells (8 and 18 proteins up- or down-regulated, respectively) (**Supp Fig 2B-C**). Additionally, gene set enrichment analysis of upregulated or downregulated factors using curated “Hallmark” pathways did not show an enrichment for proteins involved in translation in either the transcriptional or proteomic data sets (**Supp Fig 2D-E**)^79^. Taken together, these results indicate that ATL2 depletion dysregulates expression of selected cellular genes at the mRNA and protein levels but is not associated with a general shutoff of host translation.

To determine if ATL2 selectively modulates viral protein synthesis or stability, we employed the plasmid-induced replication organelle (pIRO) system, which expresses flavivirus sub-genomic RNA under the control of an exogenous T7 promoter. Upon transfection of the DENV (pIRO-D) or ZIKV (pIRO-Z) pIRO constructs into Huh7-Lunet cells expressing a cytosolic T7 promoter (Lunet-T7 cells), viral nonstructural proteins are constitutively expressed, leading to the formation of vROs (**Supp Fig 2F)**^63,64^. Thus, this system allows for specific evaluation of viral protein production and vRO formation outside the context of virus replication. To determine the impact of ATL2 KD on viral protein production, Lunet-T7 cells were transduced with lentiviruses expressing ATL2 or NT shRNA, followed by transfection with either pIRO-D or pIRO-Z. Immunofluorescence staining for viral NS3 and imaging of transfected cells showed that ATL2 depletion did not alter the transfection efficiency of pIRO-D or pIRO-Z (**Supp Fig 2G-H**). Western blot analysis showed that production of viral non-structural proteins was not significantly different between control and ATL2-depleted cells, indicating that ATL2 depletion does not specifically impair viral polyprotein production (**Fig 2E-H**). To determine if ATL2 depletion altered vRO biogenesis in the absence of viral replication, we evaluated pIRO-transfected NT and ATL2 KD cells by TEM (**Fig 2I-P**). Paralleling results from infected cells, ATL2 depletion in cells transfected with the dengue-specific pIRO-D caused clustering of vROs and reductions in vRO diameters compared to NT control cells (**Fig 2I-L**). Interestingly, in cells transfected with the Zika-specific pIRO-Z, vRO clustering was not observed during ATL2 depletion (**Fig 2M-P**); however, we did see a significant reduction in the number of cells containing vROs, indicating a decrease in the efficiency of vRO production in this expression system. Together, these results indicated that ATL2 depletion caused a defect in proper vRO formation even in the absence of viral genome replication, and that this defect is not due to changes in viral protein levels.

### ATL2 depletion increases interferon-stimulated gene activation in flavivirus-infected cells

A primary predicted function of vROs is to conceal viral RNA replication intermediates from sensing by cellular pattern recognition receptors (PRRs), including RIG-I and MDA5^16,20,21^. We therefore reasoned that altered vRO formation and organization in ATL2 KD cells might lead to increased exposure of viral RNA to PRRs. To test this, we evaluated the levels of innate immune genes in ATL2-depleted cells infected with flaviviruses. Interestingly, despite the observed reduction in viral replication, we observed a significant increase in interferon beta (IFN-β) and the interferon-stimulated genes (ISGs) MX1 and OAS1 in ATL2 KD cells (**Fig 3 and Supp Fig 3A-B**). Similar increases were not observed for the TNF-activated gene A20, although we noted a slight increase in A20 levels in ATL2 KD mock-infected cells, which was consistent with an enrichment for TNF signaling and inflammatory response genes observed by RNA-seq (**Fig 3D and Supp Fig 2d**). To rule out the possibility that ATL2 depletion generally primes cells for increased interferon activation, we also evaluated immune gene activation in alphavirus-infected cells. During ONNV or CHIKV infection, we observed ISG activation, but there was no significant difference between NT and ATL2 KD cells for the genes we investigated (**Supp Fig 3C-F**), further suggesting a specific mechanism of ATL2 in flavivirus-induced innate immune responses to infection. These data showed that ATL2 depletion leads to increased immune activation specifically in flavivirus-infected cells, possibly contributing to the observed decrease in flavivirus replication observed in ATL2 KD cells.

**Fig 3.**
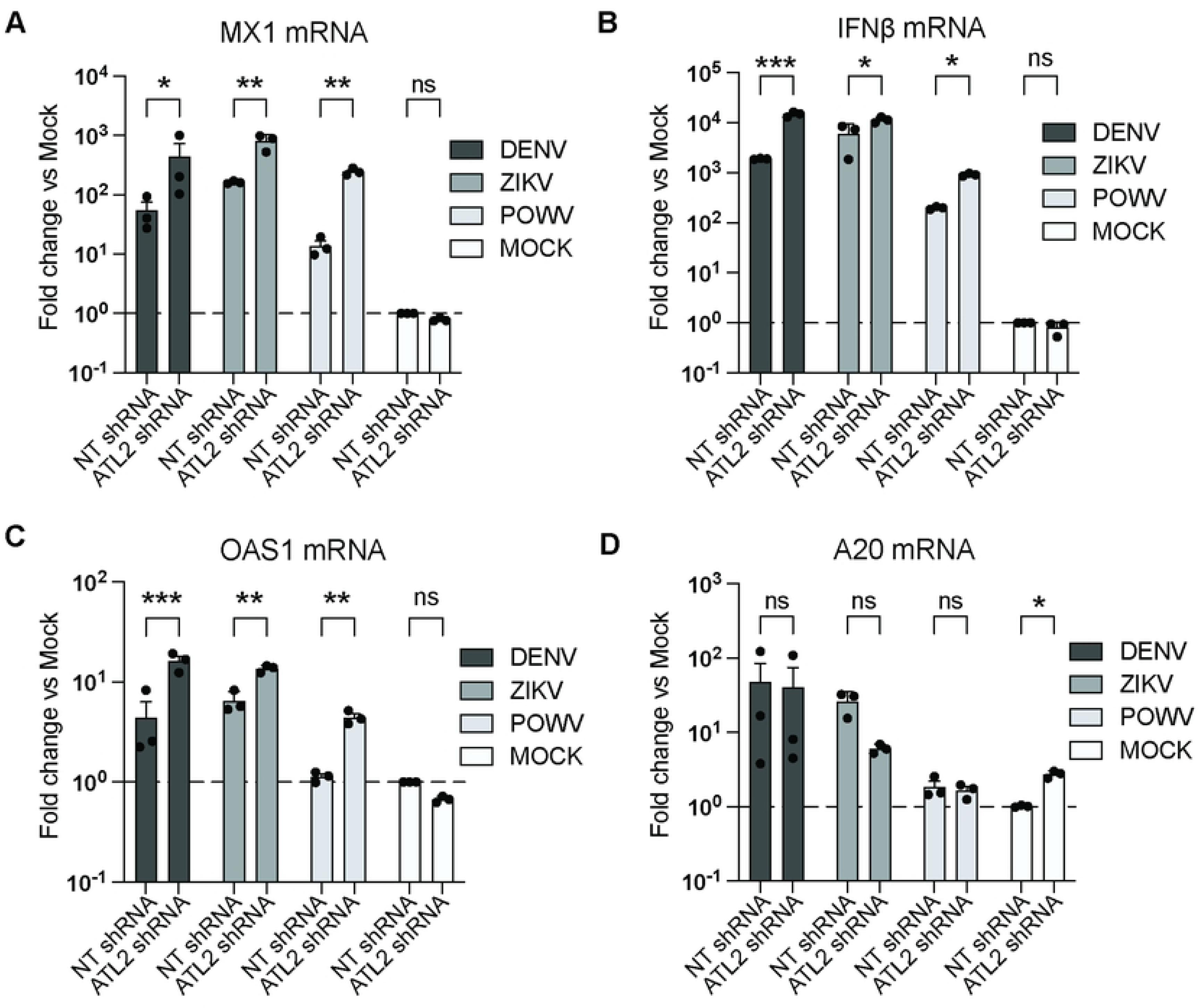
ATL2 KD alters immune gene activation in flavivirus-infected cells. Cells were transduced with constructs encoding for shRNAs directed against ATL2 or a NT shRNA. 48 hr after transduction, cells were infected with DENV, ZIKV, POWV or mock-infected for 24 hr, followed by evaluation of mRNA levels of the indicated genes by RT-qPCR. Graphs show the mean fold change in each transcript level compared to mock-infected NT shRNA cells. Statistical significance was determined using one-way ANOVA with Dunnett’s multiple comparison analysis. *P < 0.05, **P < 0.01, ***P < 0.001.

### ATL2 is enriched in cellular regions containing vROs

We next aimed to determine if ATL2 was associated with sites of flavivirus replication. A549 cells expressing HA-tagged ATL2 (HA-ATL2) were infected with DENV or ZIKV, or mock-infected, and the localization of ATL2 and viral proteins/dsRNA was evaluated by immunofluorescence and confocal microscopy. In uninfected cells, ATL2 fluorescence signal was spread in a reticular pattern throughout the cytoplasm, typical of ER localization, as previously reported (**Fig 4A**)^37,80^. However, in both ZIKV- and DENV-infected cells, ATL2 signal was enriched at sites containing both dsRNA and NS3 signal (**Fig 4A-B and Supp Fig 4A-B**). To further confirm the proximity of ATL2 to sites of virus replication, we used proximity ligation assays (PLA). This technique determines whether two proteins are associated within cells (at a maximum distance of 40 nm) and further permits the localization of intracellular protein complexes by confocal microscopy. PLAs were performed in DENV-, ZIKV- or mock-infected cells expressing HA-ATL2, evaluating the proximity of HA-ATL2 to viral nonstructural protein 5 (NS5, the viral RNA-dependent RNA polymerase). As an additional specificity control, we also evaluated PLA signal in cells expressing untagged ATL2. In HA-ATL2 cells infected with either DENV or ZIKV we observed a significant increase in PLA signal for HA-ATL2:NS5, compared to mock-infected or untagged ATL2 (**Fig 4C-D and Supp Fig 4C-D**). Since cytosolic NS5 is primarily localized to vROs, these results indicate that ATL2 is enriched in cellular regions proximal to vROs^17^.

**Fig 4.**
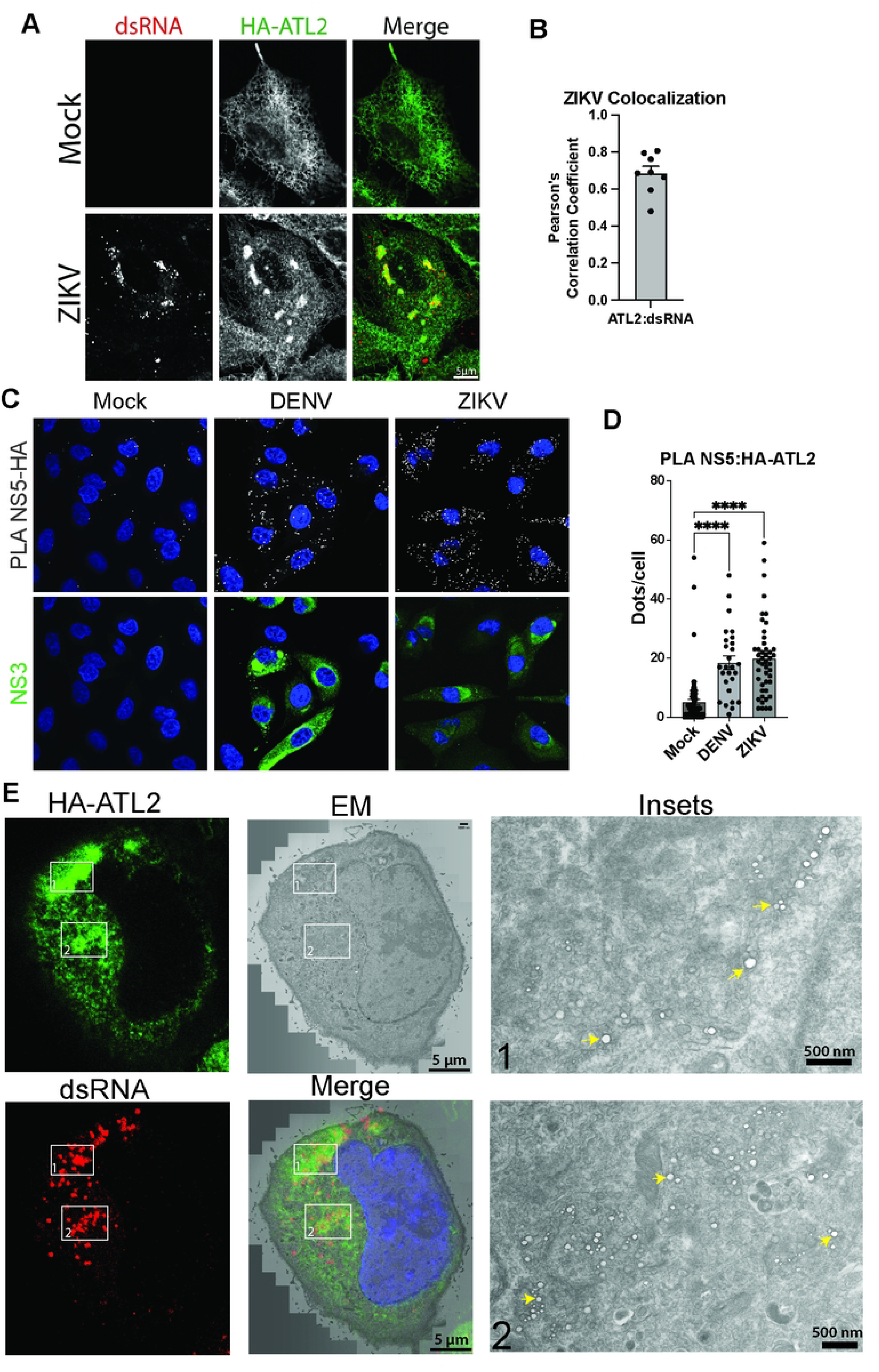
ATL2 colocalizes with sites of vRO formation during flavivirus infection. **A-D,** A549 cells expressing HA-tagged ATL2 were infected with DENV or ZIKV. **A,** 24 hr after ZIKV infection, cells were fixed and stained for dsRNA and HA using specific antibodies. Panel shows representative images for mock-infected and ZIKV-infected cells. **B,** Pearson’s correlation coefficients for the indicated fluorescent signals in the displayed images are shown in the merge channel. Graph shows the average Pearson’s correlation coefficients for the indicated fluorescence signals for ≥8 images (n≥30 cells per condition). **C-D,** 24 hr after infection with ZIKV, or 48 hr after infection with DENV, cells were fixed and subjected to proximity ligation assays (PLA) using anti-HA and anti-NS5 antibodies to detect HA-ALT2:NS5 complexes, and immunostained for NS3 viral protein to identify infected cells. Cells were imaged using confocal microscopy. Representative images are shown. Scale bar = 20 µm. **D,** Quantification of PLA dot abundance for HA-ATL2:NS5 interactions in uninfected and infected cells from two independent experiments. **E**, A549 cells expressing HA-tagged ATL2 were grown in MatTek gridded cell culture dishes and infected with ZIKV. After 48 hr, cells were fixed with paraformaldehyde and glutaraldehyde, followed by staining for dsRNA and HA using specific antibodies and reversible saponin permeabilization. Cells were imaged by confocal microscopy, then immediately embedded and processed for EM imaging using MatTek grid numbers as a reference for areas of interest. Correlation of light microscopy and EM images was done with Fiji and Adobe Photoshop software. Insets are a magnification of the boxed areas. Several individual vROs are marked with yellow arrows in the inset.

To demonstrate that areas enriched for ATL2 fluorescence signal contain vROs, we employed correlative light and electron microscopy with immunofluorescence (immuno-CLEM), which combines confocal imaging with TEM^61,62,81^. Cells expressing HA-ATL2 were infected with ZIKV for 48 hr, fixed, immunostained, and imaged using confocal microscopy, followed by processing for TEM imaging. Overlaying confocal and TEM images, we found that for ZIKV-infected cells, regions enriched for ATL2 and dsRNA signal correlated well with regions containing vROs (**Fig 4E**). Indeed, vROs were only observed in areas of enriched ATL2 signal. These results confirmed that ATL2 is present at the sites of vRO formation during flavivirus infection, consistent with a direct role in vRO biogenesis.

### ATL2 membrane tethering is critical for flavivirus replication

Atlastin-mediated membrane fusion has several well-characterized steps: following GTP binding, ATL2 monomers dimerize across membranes (loose membrane tethering), with a subsequent conformational change bringing membranes into closer association (tight tethering) and leading to eventual membrane fusion. To determine the specific function of ATL2 that facilitates flavivirus infection, we leveraged previously defined ATL2 mutations that block specific steps of this process: K107A (blocks dimerization), P371G/K372E (blocks tight tethering and fusion), and Δ524-583 (blocks fusion) (**Fig 5A-B**)^37,40,82–90^. ATL2 mutants were expressed in control or ATL2-knockout (ATL2-KO) cells (see Methods and **Supp Fig 5A**), followed by flavivirus infection and evaluation of viral replication **(Supp Fig 5B**).

**Fig 5.**
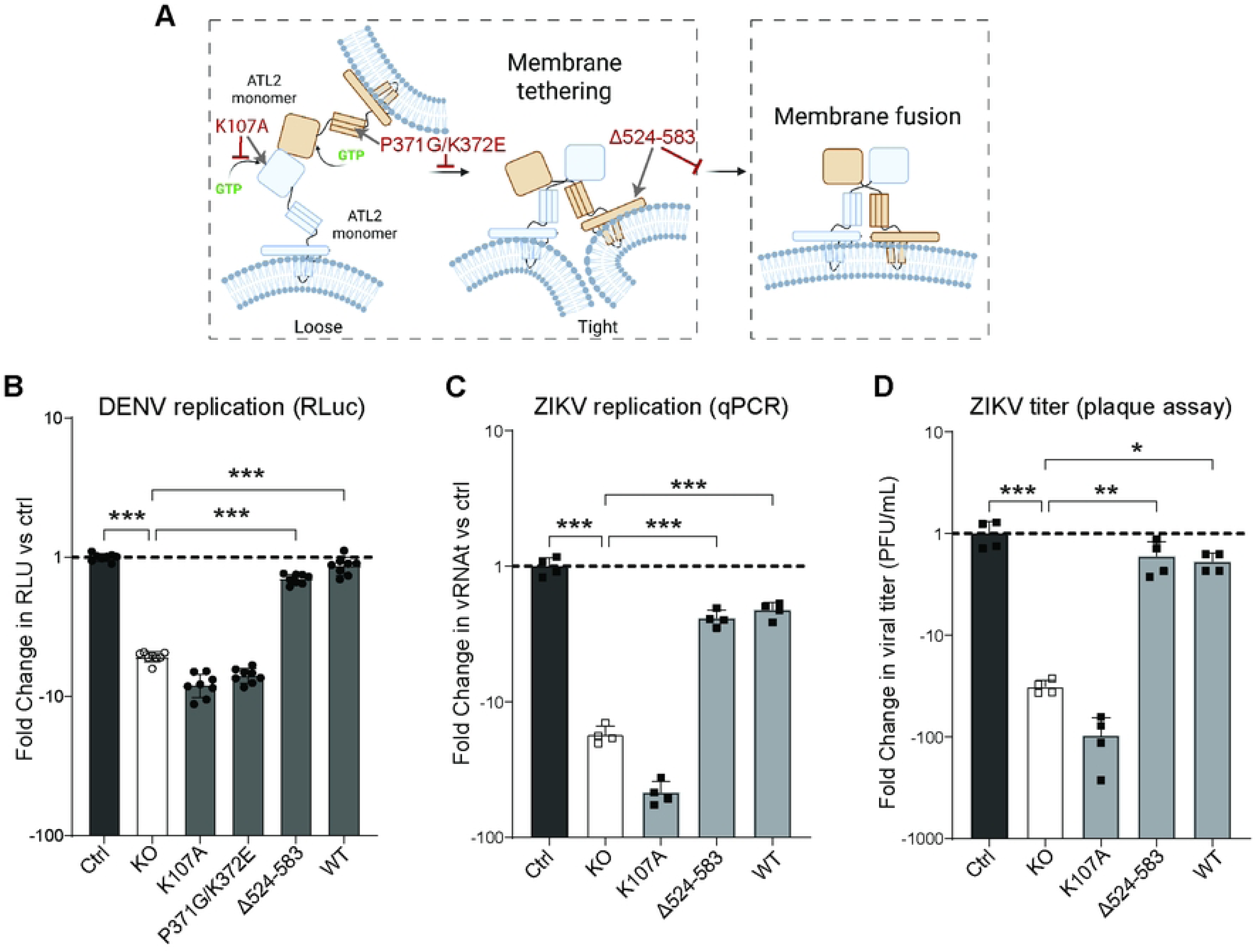
Membrane tethering-competent ATL2 variants rescue flavivirus replication in ATL2-KO cells. **A**, Schematic of the membrane tethering and fusion mechanism of ATL2, showing ATL2 mutation sites (grey arrows) and effects on tethering and/or fusion (red inhibitors). **B-C,** Graphs show the average fold change in DENV (**B**) or ZIKV (**C**) replication in cells expressing the indicated ATL2 mutants. **D**, Graph shows the average fold change in ZIKV titers (PFU/mL) harvested from cells expressing the indicated ATL2 mutants (measured by plaque assay). For all graphs, results were normalized to viral replication in control cells expressing an empty lentiviral vector (left bar in each graph). All other bars display data from ATL2-KO cells expressing the indicated mutants. Statistical significance was determined by one-way ANOVA with Dunnett’s multiple comparison analysis. *P < 0.05, **P < 0.01, ***P < 0.001.

In cells infected with *Renilla* luciferase-expressing DENV (DV2-R2A), expression of full-length wildtype (WT) ATL2 restored DV-R2A replication in ATL-KO cells to levels comparable to control cells. Importantly, overexpression of WT ATL2 did not significantly alter DV-R2A replication in control cells **(Supp Fig 5C**). Expression of the GTP-binding mutant ATL2 K107A, which inhibits ATL2 dimer formation, was unable to restore DV-R2A replication in ATL2-KO cells, indicating that ATL2 GTP nucleotide binding is required for flavivirus replication (**Fig 5B and Supp Fig 5D**)^37,88^. Similarly, expression of a double mutant, ATL2 P371G/K372E, which is capable only of loose membrane tethering, did not rescue virus replication^82,88^. However, expression of a mutant which is missing the C-terminal amphipathic helix of ATL2 (ATL2 Δ524-583), required for membrane fusion but not for membrane tethering, restored DENV replication in ATL2-KO cells to levels comparable with cells expressing WT ATL2^40,82,90,91^. To confirm conservation between flaviviruses, these experiments were repeated with ZIKV. Similar to DENV-infected cells, only WT ATL2 and the Δ524-583 mutant showed significant rescue of virus replication in ATL2-KO cells (**Fig 5C-D and Supp Fig 5E-F**). Taken together, these results indicate that the primary biological function of ATL2 during flavivirus replication is membrane tethering, rather than the canonical membrane fusion typically associated with atlastin proteins.

Building on these results, we next investigated the ATL2 N-terminal hypervariable domain (HVD), which has been linked to membrane tethering. Specifically, recent reports suggest that the HVD of atlastin-1 facilitates the formation of homo-oligomeric structures that are proposed to build long membrane tethers^92^. We had previously shown that expression of an ATL2 N-terminal deletion mutant (ATL2-Δ2-65) did not result in the rescue of flavivirus replication in ATL2-KO cells^35^. To further investigate whether the ATL2 HVD functions to facilitate membrane tethering, we investigated the effect of overexpressing this domain on flavivirus infection, which based on previous data is predicted to compete for binding sites and disrupt tethering^92^. In cells overexpressing exogenous ATL2 HVD, we observed reduced DENV replication, approaching the level of reduction seen during ATL2 depletion (**Supp Fig 5G-H**). These results confirm the importance of the ATL2 HVD in viral replication and further implicate ATL2-mediated membrane tethering as the critical function of ATL2 during infection.

### ATL2 function during flavivirus infection is not regulated by phosphorylation or alternative splicing

Atlastin-mediated membrane tethering and fusion have been primarily characterized using *in vitro* membrane fusion assays; as such, mechanisms of modulation in cells are still unclear. We therefore aimed to determine how ATL2 membrane tethering is regulated in flavivirus-infected cells. Based on our results indicating the importance of the ATL2 HVD in flavivirus infection, as well as previous studies showing hyperphosphorylation of the HVD, we hypothesized that HVD phosphorylation may regulate ATL2 membrane tethering function in flavivirus replication^82,92,93^. To test this, each serine and threonine residue in the ATL2 HVD was mutated to alanine to ablate potential phosphorylation sites (**Supp Fig 6A**), and mutants were expressed in ATL2-KO or control cells, followed by infection with DV-R2A^94^.

Surprisingly, expression of ATL2 phosphorylation mutants completely restored DENV replication in ATL2-KO cells, indicating that the ATL2 function relevant for flavivirus replication is not regulated by phosphorylation (**Supp Fig 6B-C**).

We next explored the role of alternative splicing in regulating ATL2 activity. There are two primary isoforms of ATL2, as well as three alternatively-spliced isoforms which have been experimentally observed and seven additional isoforms that have been defined computationally^95,96^. To test if ATL2 isoforms are functional in restoring flavivirus replication in ATL2-KO cells (in which production of all ATL2 isoforms was ablated), five representative isoforms were expressed in ATL2-KO or control cells, followed by infection with DENV and evaluation of virus replication (**Supp Fig 6D-F**). The only isoform that failed to restore viral replication was ATL2-3 (UniProt identifier Q8NHH9-3), which lacks the GTPase active site (**Supp Fig 6D**)^96^. While we did not test all possible isoforms, the isoforms tested were representative of other predicted variants, indicating that alternative splicing of ATL2 likely does not have a significant impact on flavivirus replication.

### ATL2 activity can be targeted with a peptide inhibitor to reduce flavivirus replication

Having determined the significance of ATL2 in flavivirus replication, we next aimed to leverage this knowledge to inhibit viral replication. A previous report identified an autoinhibitory amino acid sequence within the ATL2 C-terminus that can be used as an exogenous peptide inhibitor of ATL2 function (**Fig 6A**)^93^. To test if this peptide could be used in cells to limit ATL2 activity and inhibit flavivirus replication, we added a cell-penetrating HIV-TAT sequence to this peptide (pepTAT-A2), applied it to A549 cells, infected these cells with DENV, and evaluated viral replication using immunofluorescence microscopy and plaque assay^97^. Strikingly, the percentage of virus-infected cells, as well as viral titers from infected cells, were significantly decreased upon treatment with TAT-ATL2 compared to cells treated with a pepTAT control peptide (**Fig 6B-D**). To confirm the specificity of the peptide for ATL2, we also compared results to cells treated with a modified peptide containing three charge reversals (pepTAT-A2-KKE), which exhibits decreased interactions with ATL2 and approximately 50 percent less inhibitory activity *in vitro*^93^. Treatment with the pepTAT-A2-KKE peptide still led to moderate reductions in virus infection, but its antiviral activity was markedly attenuated (**Fig 6B-D**). As a further confirmation of specificity, a peptide inhibitor without the cell penetrating TAT sequence did not affect virus replication (**Supp Fig 7A**). None of the peptide treatments had a significant impact upon cell viability (**Supp Fig 7B**). These results strongly suggest that the effect of pepTAT-A2 on viral replication is specific to its inhibition of ATL2 function.

**Fig 6.**
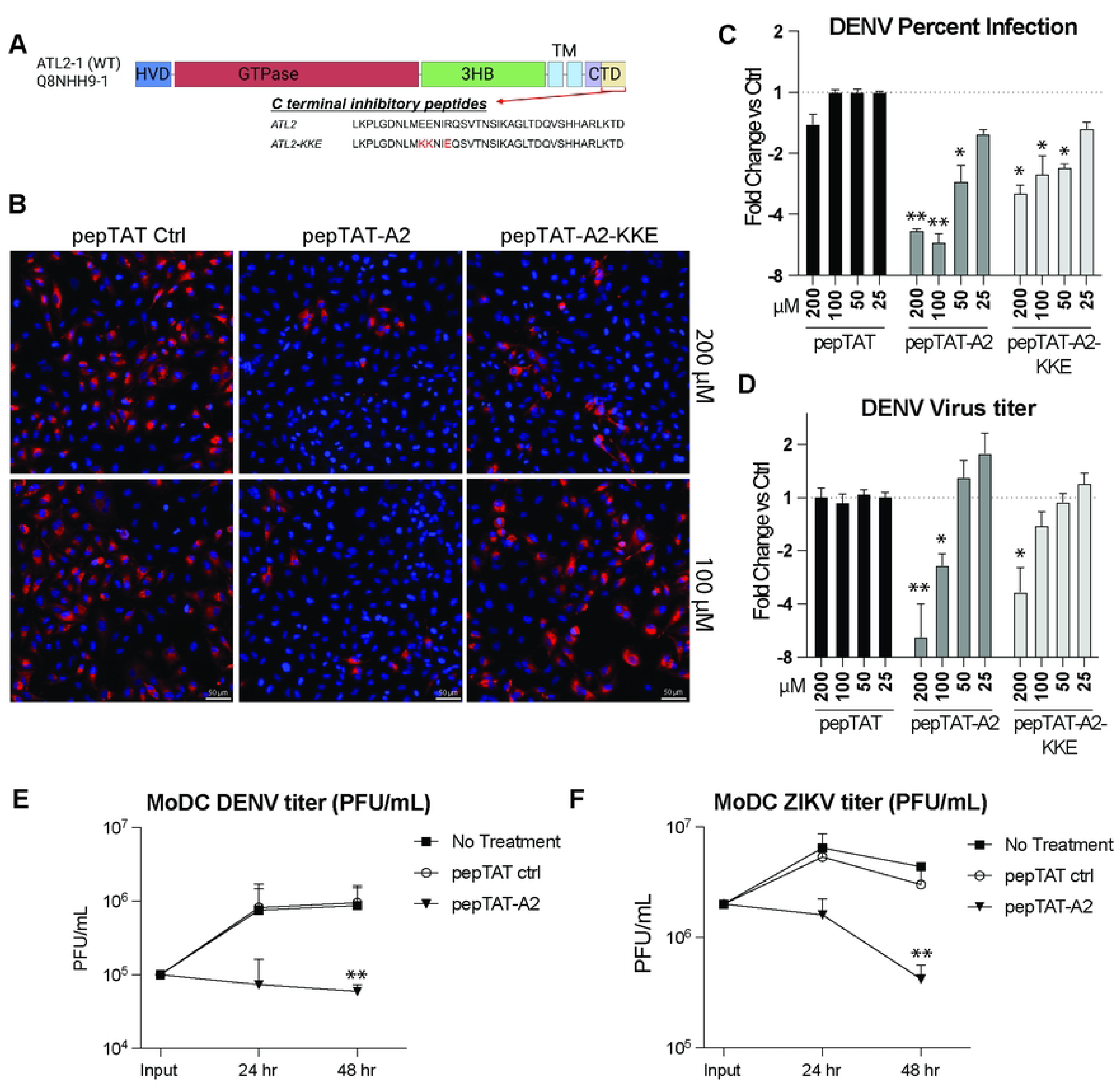
ATL2 inhibitory peptides limit flavivirus replication in immortalized and primary cells. **A,** Schematic representation of the primary ATL2 isoform (ATL2-1, UniProt identifier Q8NHH9-1), highlighting the C-terminal autoinhibitory peptide. **B-D,** A549 cells were treated with the indicated peptide at the indicated concentrations and infected with DENV (MOI: 1) for 48 h. **B,** Cells were fixed with PFA and viewed by immunofluorescence microscopy using antibodies specific for the viral NS3 protein. Images are representative of 4 independent experiments. **C,** Graph shows the average fold change in percentage of infection for each peptide at the indicated concentration. **D,** Graph shows the average fold change in virus titer from supernatants of cells treated with each peptide at the indicated concentration. **E-F,** MoDC cells were treated with the indicated peptide at 100 µm and infected with either DENV (**E**) or ZIKV (**F**). Graphs show the average PFU/ml for supernatants at each time point. Graphs represent duplicate samples from 3 independent donors. Statistical significance was determined using one-way ANOVA with Dunnett’s multiple comparison analysis. Significance shown for each sample is compared to pepTAT control samples. *P < 0.05, **P < 0.01, ***P < 0.001.

To confirm our observations in a more biologically relevant system, we evaluated the effect of peptide treatment in primary human monocyte-derived dendritic cells (moDCs). These cells are frequently among the first sites of viral replication upon natural infection and are important for establishing and spreading flavivirus infections within human hosts^98–100^. MoDCs differentiated from human peripheral blood mononuclear cells (PBMCs) were incubated with the indicated peptides and infected with either DENV or ZIKV^73–75^. Productive infection in moDCs was confirmed by verifying a significant loss of viral titers upon addition of a pan-flavivirus polymerase inhibitor (NITD008)^101,102^. Paralleling results from A549 cells, treatment with pepTAT-A2 significantly reduced titers and intracellular viral RNA levels for both viruses, as compared to the pepTAT control peptide, without impacting cell viability (**Fig 6E-F and Supp Fig 7C-E**). Thus, across both immortalized and primary human cells, our results demonstrate that ATL2 function can be targeted to inhibit flavivirus infection.

## Discussion

Defining the membrane reorganization mechanisms that govern vRO biogenesis and RNA trafficking in infected cells is critical to understanding flavivirus infection. Here, we present evidence that the spatial organization of flavivirus genome replication is mediated by the membrane tethering capability of the host factor ATL2. Depletion of ATL2 in flavivirus-infected cells impeded proper vRO formation and reduced virus replication, which correlated with a robust increase in cellular immune activation. While the membrane tethering activity of ATL2 was required for productive flavivirus infection, its membrane fusion activity proved dispensable. Based on these findings, we propose a model in which the organization and distribution of flavivirus vROs in the cytoplasm of infected cells is mediated by homotypic ER membrane tethers (**Fig 7**). This organization allows newly synthesized viral genomes egressing from existing vROs to form *de novo* vROs on juxtaposed ER membranes. The reversible quality of ER tethers then allows the dispersion of vROs throughout the cell as the ER is dynamically rearranged^40,82,86,87,103,104^. Without ER membrane tethers, viral RNA remains confined to *cis*-ER membranes, resulting in the accumulation of vROs and viral RNA at the initial replication site. Our data also suggest that ER membrane tethers may have an immune-protective role, potentially limiting access of pattern recognition receptors (PRRs) to viral RNA intermediates. Together, these findings indicate that ER-to-ER contacts are needed to assemble specific membrane microdomains that are critical for sustaining flavivirus replication.

**Fig 7.**
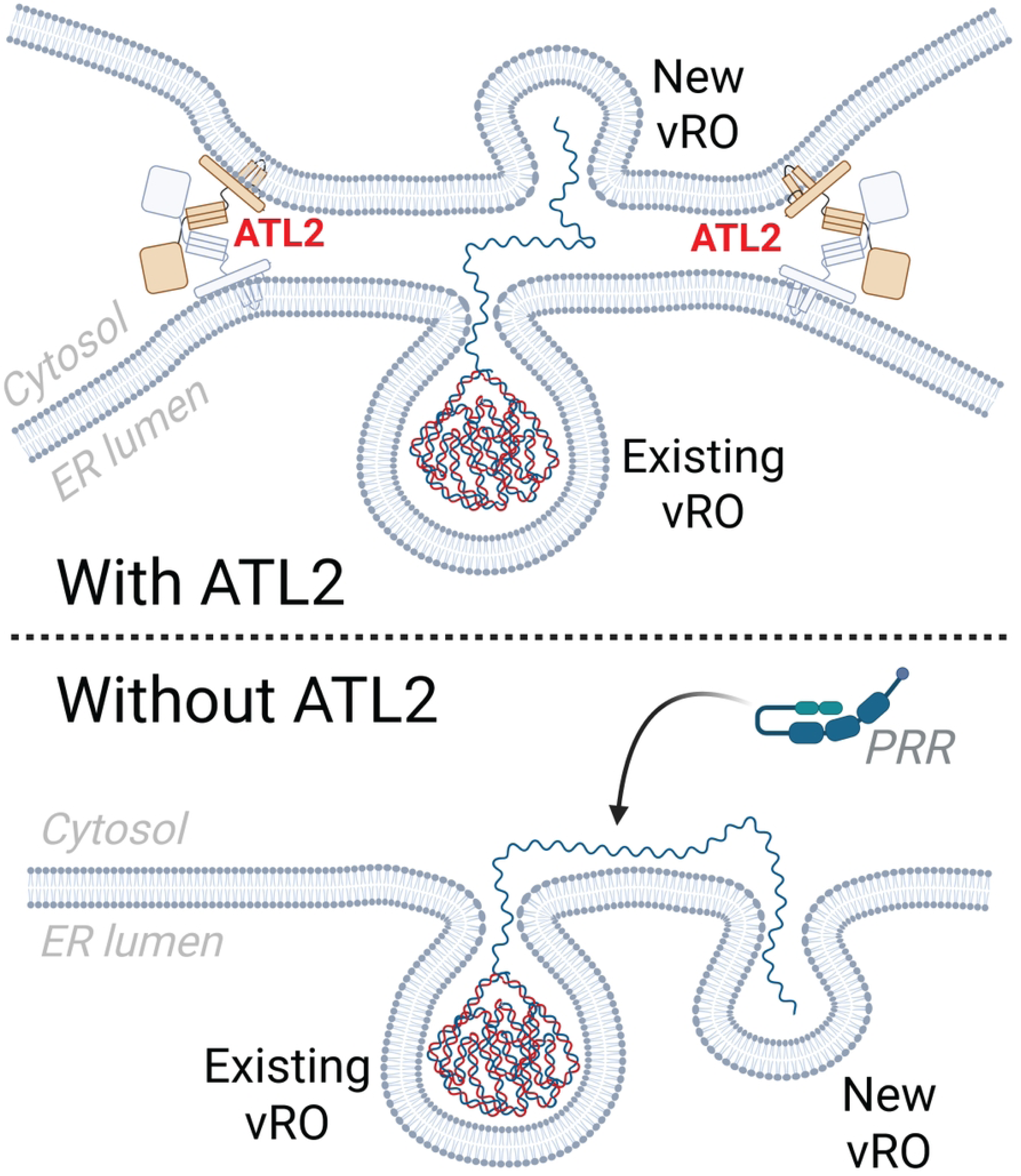
Model: ATL2 membrane tethering is critical for the spatial organization of flavivirus genome replication. ATL2 tethering dimers bring ER membranes into close contact, facilitating the trafficking of newly synthesized viral genomes from existing vROs into vROs forming on juxtaposed membranes (top). This close connection forms a microdomain which brings together the necessary factors for nascent vRO formation and shields the cytosol-exposed viral RNA from innate immune sensors. Because membrane tethering is reversible, this also permits the expansion of viral genome replication away from sites of existing vRO biogenesis. In the absence of ATL2 membrane tethering (bottom), viral genomes extruded from the vRO are exposed to the cytosol and more likely to form new vROs on the same membrane, leading to the concentration of sites of vRO biogenesis and increased recognition by pattern recognition receptors (PRRs) in the cytosol.

Our evaluation of DENV-infected cells by electron tomography (ET) revealed numerous membrane contact sites between vROs and juxtaposed membranes. These results parallel previous findings for DENV and other flaviviruses, including recent observations for tick-borne encephalitis virus (TBEV) using *in situ* cryo-ET^17,18,26^. In both DENV- and TBEV-infected cells, smaller vROs were often found in ER membranes tightly juxtaposed to other ER membranes, potentially representing intermediates of vRO formation before genome replication is established. These observations suggest that membrane contacts also contribute to proper vRO maturation. This is supported by our data showing decreased genome replication and enrichment of smaller vROs in ATL2-depleted cells. Furthermore, ET has also demonstrated the tight membrane contacts that occur at the viral assembly site, where the juxtaposition of ER membranes enables direct transfer of genomic RNA from the vRO into a budding virion^17–19,26,105^.

This arrangement likely improves the efficiency of virus production by concentrating required viral assembly components in one location. In ATL2-depleted cells, we observed very few virions by electron microscopy and, indeed, significantly decreased viral titers in supernatants, suggesting that ATL2 may also be required to establish the membrane contacts needed for virion assembly. Taken together with prior studies, our results support the importance of ER-to-ER membrane contacts for vRO maturation and perhaps for virion budding.

In addition to reducing viral replication, ATL2 depletion and the resulting disruption of ER membrane tethers increased cellular immune activation. A key predicted function of vROs is to sequester replication machinery from the cytosol, thereby shielding replication intermediates from immune receptors such as RIG-I and MDA5^16,20,21^. Increased immune activation in ATL2-depleted, flavivirus-infected cells suggests that these receptors have greater access to viral genomic material. This could result from aberrant release of dsRNA from vROs, or increased exposure of single-stranded (ssRNA) or double-stranded RNA moieties. Since vROs still assemble without ATL2, immune activation is more likely due to increased recognition of ssRNA, potentially due to trafficking disruptions once newly synthesized ssRNA exits the vRO. In addition to detecting dsRNA, RIG-I can recognize 5’-triphosphate ends on ssRNA^106^.

Our data suggest that membrane connections may create a cellular microdomain that restricts PRR access, allowing viral RNA to traffic between membranes undetected.

Alternatively, changes in vRO morphology and distribution in ATL2-depleted cells may reduce capping efficiency, leading to more 5’-triphosphate ends detectable by RIG-I. We also noted that in ATL2-depleted cells, there was a slight enrichment for inflammatory and TNF gene upregulation, which could impact immune activation and flavivirus infection. However, previous studies have demonstrated that although TNF and inflammation play a role in DENV disease severity in patients, treatment with TNF does not directly impact viral replication^107–109^. Additionally, we show that ATL2 depletion does not impact immune activation or viral replication in alphavirus-infected cells, indicating that the immune activation and reduction in viral replication observed in flavivirus infection is specific and not a result of general inflammatory changes from ATL2 depletion. Clarifying the mechanisms of immune activation in the absence of ATL2 will be the focus of future studies.

Our data show that ER membrane tethering has a biological role during infection, which may extend to other aspects of cellular homeostasis. Atlastin proteins are best known for mediating ER tubule fusion, yet the low intrinsic fusion activity of the predominant ATL2 isoform and *in vitro* evidence of reversible, ATL2-mediated liposome tethering support a role for ATL2 membrane tethering in regulating cellular functions.

However, to our knowledge, only one other study has shown a biological relevance for atlastin tethering, specifically showing that the membrane tethering function of atlastin-1 facilitates COPII-coated vesicle trafficking in neurons^39^. However, these conclusions were predicated upon the use of a fusion mutant which has been shown to also display tethering deficiencies, and this work has not yet been replicated for ATL2^40,82,89^. Building on our results implicating ATL2 in viral RNA trafficking, we speculate that ATL2 may also mediate trafficking of cellular factors. Since transcriptomic and proteomic analyses of ATL2-depleted cells did not reveal broad translational changes, it is less likely that mRNA trafficking is altered; however, it is possible that specific protein or lipid distributions are altered. Alternatively, ATL2-mediated tethers may have other functions in maintaining ER structure and homeostasis. Understanding how ATL2 fusion and tethering are regulated will be key to determining how these activities contribute to cellular functions.

Recent studies examining atlastin protein fusion and tethering activities have identified N- and C-terminal elements that regulate ATL2^42,82,92,93^. Here, we demonstrate that providing excess amounts of either the ATL2 N-terminal HVD or the ATL2-1 C-terminal autoregulatory sequence modulates ATL2 activity and reduces viral replication in both immortalized and primary cells. The impact of ATL2 modulation upon flavivirus replication and specifically vRO formation highlights intriguing avenues for future research.

Conservation of vRO morphology among flaviviruses makes defining the structure and function of vROs highly relevant for both fundamental understanding of flavivirus biology and for identifying potential therapeutic targets. Indeed, several recent studies have illustrated the antiviral potential of drugs which disrupt vRO assembly and maintenance^110–113^. However, little is known about the mechanisms of vRO biogenesis and, in particular, the importance of host factors in the formation and function of these complexes. Our data provide evidence that a mechanistic understanding of host factor function in flavivirus infection can inform target selection for antiviral development. Defining how host factors coordinate vRO formation could expand the field of host-targeted antivirals, which – due to the common mechanisms among flavivirus infections and the theoretical evolutionary barrier presented by host-targeted therapies – possess great potential for broad-spectrum activity^10,11^.

In summary, we show that ATL2-mediated membrane tethering serves as a critical link in the chain of events that facilitate a productive flavivirus infection. The role of ATL2 tethering in facilitating infection is conserved for multiple flaviviruses, representing a common host mechanism in flavivirus replication. This knowledge reinforces the importance of spatial coordination of viral processes during infection, sheds new light upon molecular mechanisms of ER regulation, and illuminates a specific and conserved relationship between flaviviruses and their hosts.

## Acknowledgments

This work was supported by the Emory University Integrated Cellular Imaging Core Facility (RRID:SCR_023534). Research reported in this publication was supported by the Office of The Director of the National Institutes of Health under Award Number S10OD028673 (RRID:SCR_023534). This study was also supported by Emory University Robert P. Apkarian Integrated Electron Microscopy Core Facility (RRID: SCR_023537), which is subsidized by the Emory University School of Medicine and Emory College of Arts and Sciences. Additional support was provided by the Georgia Clinical & Translational Science Alliance of the National Institutes of Health under award number UL1TR000454. Electron microscopy data described here were collected on a JEOL JEM-1400, 120kV TEM supported by the National Institutes of Health Grant S10 RR025679. This study was also supported in part by the Emory Integrated Genomics Core (EIGC; RRID:SCR_023529), which is subsidized by the Emory University School of Medicine and is one of the Emory Integrated Core Facilities. Additional support was provided by the Georgia Clinical & Translational Science Alliance of the National Institutes of Health under Award Number UL1TR002378.

Work in Dr. Christopher Neufeldt’s laboratory was supported by The National Institute of Allergy and Infectious Diseases (1R01AI185849-01). Work in Dr. Pietro Scaturro’s laboratory was supported by the Free and Hanseatic City of Hamburg and the German Research Foundation (DFG, Deutsche Forschungsgemeinschaft) under Grant 499961789, as well as the Federal Ministry for Education and Research (BMBF, Bundesministerium für Bildung und Forschung) under grant 13GW0622. PS is associated with the DFG Collaborative Research Center (CRC) 1648 (SFB 1648/1 2024—512741711). We also acknowledge the support of the Microbiology and Molecular Genetics PhD Program at Emory University.

The content of this work is solely the responsibility of the authors and does not necessarily reflect the official views of the National Institutes of Health.

## Author contributions

Conceptualization, JEO and CJN; Investigation, JEO, CLB, QW, ACV, JWVV, LA, OU, MG, CDS, PS, CJN; Writing – Original Draft, JEO and CJN; Writing – Review & Editing, JEO, CDS, LCC, PS, MSS, CJN; Funding Acquisition, LCC, PS, MSS, and CJN.

